# Identifying and ranking species that need urgent management action to achieve Target 4 of the Global Biodiversity Framework

**DOI:** 10.64898/2026.04.11.717432

**Authors:** H. Resit Akcakaya, Natasha L. M. Mannion, Jonah Morreale, Domitilla Raimondo, Michael Hoffmann, Stuart H. M. Butchart, Louise Mair, Francesca A. Ridley, Malin Rivers, Caitlin Brant, Michael Clifford, Megan Joyce, Kira Mileham, Celia Nova Felicity, Mirza Kusrini, Sunarto, Johnpaul Houston, Nunia Thomas, Simon T. Maddock, José F. González-Maya, Kostas Triantis, Dimitrios Vavylis, Konstantina Spiliopoulou, Indira A. Gamatis, Bello A. Danmallam, Samuel T. Ivande, Shiiwua A. Manu, Stella Egbe, Joseph D. Onoja, Carolina Castellanos-Castro, Cristina Lopez-Gallego, Barney Long, Philip J. K. McGowan

## Abstract

Target 4 of the Kunming–Montreal Global Biodiversity Framework (KMGBF) calls for urgent management actions to halt human-induced extinctions and enable species recovery. However, most Parties face substantial challenges in determining which species require urgent management actions. Here, we present a transparent, standardised protocol that identifies and ranks species most likely to need urgent management actions at the national level, using globally available data from the IUCN Red List of Threatened Species. The protocol integrates four criteria aligned with Target 4: global extinction risk, rate of decline, population or range restriction, and endemism, to generate a national ranked list of species. Species scoring highly on these four criteria, and therefore most in need of urgent management action, are ranked most highly. We applied this method to all 250 countries and territories listed in the IUCN Red List and pilot-tested national rankings with participants from eight diverse countries. Across pilots, participants reported that the ranked lists were scientifically robust, time-saving, and valuable starting points for national priority-setting, while stating the importance of national context, and the need for additional technical and financial support for implementation. Our results demonstrate that a science-based approach can meaningfully support Parties in identifying species requiring urgent action under Target 4, in a standardised way. With 2030 approaching rapidly, this protocol provides an immediate, practical tool to accelerate progress toward halting extinctions and advancing species recovery.

## 1. Introduction

The world is facing a global biodiversity crisis driven by human activity, requiring widespread transformative change (Díaz *et al*., 2019). Over 48,000 species are documented as threatened with extinction (IUCN, 2025); extrapolations suggest the true figure may exceed 1 million species (IPBES, 2019). In response to declines in biodiversity, governments have adopted international agreements including the Convention on Biological Diversity (CBD) (United Nations, 1992).

Parties to the CBD adopted the Strategic Plan for Biodiversity for 2011-2020, which included the Aichi Biodiversity Targets, with the aim of conserving and achieving sustainable use of biodiversity by 2020 (CBD, 2010). Parties did not fully achieve any of the targets (Secretariat of the CBD, 2020a, IPBES, 2019), including the target for reducing risk of species extinction (Aichi Biodiversity Target 12). Whilst there was some success in avoiding extinctions of known threatened species (Bolam *et al*., 2021), the conservation status of threatened species is not improving (Secretariat of the CBD, 2020b), and overall species are moving towards extinction at an increasing rate (IPBES, 2019), as quantified by the Red List Index (Butchart et al. 2004). The scale of the challenge and the absence of a strategic approach to identifying where action would be most impactful in reducing overall extinction risk meant that responses were often fragmented and focussed on individual, relatively high profile, species and actions (see CBD 2016a).

The successor to the CBD’s Strategic Plan for Biodiversity is the Kunming-Montreal Global Biodiversity Framework (KMGBF), which was adopted in 2022 and also contains clear aspirations for species conservation. It includes four outcome-orientated Goals to start the road to nature recovery by 2050, to be progressed towards by 23 action-orientated targets to be achieved by 2030 (CBD, 2022a). Goal A includes the ambition that “*Human induced extinction of known threatened species is halted, and, by 2050, the extinction rate and risk of all species are reduced tenfold and the abundance of native wild species is increased to healthy and resilient levels.”*

To contribute towards meeting Goal A of the KMGBF there are eight Targets specifically concerned with ‘reducing threats to biodiversity’. Of these, Targets 1-3 and 5-8 are concerned with reducing particular threats to biodiversity and are based on the five main drivers of biodiversity loss identified in the IPBES Global Assessment: changes in land and sea use, direct exploitation of organisms, climate change, pollution, and invasion of alien species (IPBES, 2019). For some species the responses implemented through these seven Targets will be insufficient to prevent their extinction and enable their recovery, and additional action will be required (Bolam *et al*., 2023). Target 4 is specifically included in the KMGBF in recognition of this, and hence actions targeted directly at specific species are needed to avoid extinction and enable recovery (McGowan *et al*., 2024).

Target 4: “*Ensure urgent management actions to halt human induced extinction of known threatened species and for the recovery and conservation of species, in particular threatened species, to significantly reduce extinction risk, as well as to maintain and restore the genetic diversity within and between populations of native, wild and domesticated species to maintain their adaptive potential, including through in situ and ex situ conservation and sustainable management practices, and effectively manage human-wildlife interactions to minimize human-wildlife conflict for coexistence*.”

One of the challenges faced by Parties in achieving Aichi Biodiversity Target 12 was obtaining “a clear understanding and an explicit statement of which species should be the focus of conservation action” (CBD, 2016b). Given many countries are home to hundreds of threatened species, with 22 countries reporting more than 1000 (IUCN, 2025), this represents a challenge of considerable scale. Limited capacity and resources were notable constraints to achieving the Aichi Biodiversity Targets (Peña Moreno and Romero, 2018; Secretariat of the CBD, 2020a), and developing tools to identify and prioritise species in need of focused action under Target 4 was identified as a key area of capacity development required to achieve the GBF Targets (Maggs, Slater and McGowan, 2022). With this in mind, a standard, science-based approach to identifying native wild species that could be considered in need of urgent management action may be a helpful starting point for Parties.

Here we develop and test a method for ranking species in need of urgent management actions in each country. The overriding need is for a method that is conceptually robust, straightforward to use and can be mobilised quickly to support all Parties in addressing Target 4. We propose a protocol for identifying and ranking these species, based on risk of extinction, created through an automated compilation of information available in the global IUCN Red List of Threatened Species. A national ranked list is intended as a starting point, to be considered and reviewed by that country, before adopting its list of species in need of urgent management action. We then piloted this protocol with a diverse set of eight countries, to 1) assess whether the protocol is useful to CBD Parties in achieving Target 4, 2) understand for each Party what the next steps would be to finalise a list of species in need of urgent management action, and 3) determine whether support, in addition to the ranked list, is needed by Parties to act on this list. We present the results of these pilots, illustrating the utility of this protocol for identifying species in need of urgent management action, and supporting countries to deliver on Target 4 by 2030.

## 2. Methods

### 2.1 Structure and justification of the proposed protocol

Here we focus on the following components of Target 4 of the Global Biodiversity Framework, “*Ensure urgent management actions to halt human induced extinction of known threatened species and for the recovery and conservation of species, in particular threatened species, to significantly reduce extinction risk.”* This wording essentially translates into four considerations: (i) “urgent”, (ii) “management actions”, (iii) “threatened species”, and (iv) “significantly reduce extinction risk”.

In relation to the species for which these considerations apply, *“urgent”* suggests priority for species that are fast approaching extinction, especially those that are declining rapidly. “*Management actions”* suggests priority for species that require species-specific actions, which are often those with small populations and restricted ranges. “*Threatened species*” can be interpreted as those assessed as Critically Endangered (CR), Endangered (EN) and Vulnerable (VU), which in the IUCN Red List system are those facing a high to extremely high risk of extinction in the wild; however, we also include Extinct in the Wild (EW), as these species are entirely dependent on ex-situ conservation to avoid extinction (Smith *et al*., 2023) and because Target 4 specifically references ex-situ conservation. “*Significant reduction of risk”* suggests priority on those species at the highest risk categories, because a transition from CR to EN reduces extinction risk much more than a reduction from, say, VU to Near Threatened (NT) (Butchart *et al*., 2004). In addition, our intention to help guide countries suggests a priority on species for which a country has the highest global responsibility (e.g., species endemic to the country).

Based on these considerations, we propose a protocol that prioritises species using four criteria:

(a) Extinction risk category, to include known globally threatened species, and to ensure that the most significant reductions in extinction risk are prioritised;
(b) Rate of decline, to prioritise species requiring the most urgent actions (recognising that severe population restriction may also imply urgency);
(c) Population restriction, to prioritise species whose recovery will likely require species-targeted management actions (recognising that rapid decline may also indicate the need for targeted conservation); and
(d) Proportion of the range in the focal country, to identify species for which the country has the greatest responsibility for their global conservation.

In particular, we rely on metrics related to species restriction (small population or range) to infer a need for species-specific management actions (such as conservation translocations, recovery actions, and ex situ conservation; see [Guidance Notes of CBD](https://www.cbd.int/gbf/targets/4)). Most widespread or abundant species, even if they are globally threatened, are unlikely to benefit substantially from species-specific management actions because they are not feasible to implement at sufficient scale; the threats these species face are more feasible to address through policy responses covered under the other CBD targets (e.g. regulations to control unsustainable use under Target 5). Most species-targeted management actions are more appropriate for conserving, and typically directed towards, small and restricted populations. Our goal here is not a definitive list of species that must be conserved with species-targeted actions, but rather a list of species that are most likely to need such actions. In addition, because of the automated compilation of this information from the Red List, each species is not considered in the local context of the focal country. Thus, the list is intended for each Party to consider as a starting point for identifying species that are a priority for management action, with the expectation that it will be modified with appropriate additions and deletions, which we discuss below.

### 2.2 IUCN global Red List data

The proposed protocol applies these four criteria to information available in the global IUCN Red List of Threatened Species, described in section 2.3. The IUCN Red List is the largest global inventory of the extinction risk and conservation status of animal, fungus, and plant species, which classifies species into nine Red List Categories according to global extinction risk (IUCN, 2025).

Species are evaluated on five criteria (A-E), based on geographic range, population size, population decline or increase, and extinction probability analysis, to determine their appropriate Red List Category (IUCN, 2012). Currently over 172,600 species have global Red List assessments (IUCN, 2025).

Here we considered all taxonomic groups with species assessed for the Red List, including both groups in which all species have been assessed (e.g. mammals, birds, cycads) and those in which only a subset have been assessed (e.g. fungi, invertebrates, plants, and freshwater fishes). We acknowledge that assessments are often biased towards charismatic or better-known species, which will be reflected in the ranked lists of species produced by the protocol. We have compiled information including Red List category, Red List criteria, and other fields for each species from the most recent version of the IUCN global Red List (version 2026-1), accessed via the IUCN Red List API version 4. The reproducible pipeline including data retrieval via the Red List API and all data processing is available online (github.com/JonahMorreale/RedList_Target4Species).

### 2.3 Implementation of the proposed protocol

The output of the proposed protocol is a ranked list of species, composed of two parts. The first part of the list, i.e. the highest ranked, includes species that score on all four criteria (Priority 1); the second part, i.e. lower ranked species, includes species that score on three of the four criteria (Priority 2). Both parts of the list capture all threatened species, included on the IUCN global Red List, that are native to the focal country (including by reintroduction or assisted colonisation). Within each part of the list, species are ranked according to a priority score, with species most at risk of extinction without urgent management actions ranked as the highest priority. Additionally, EW species are placed unranked at the top of the combined lists.

The below protocol was implemented in R 4.5.2 (R Core Team, 2025) utilizing the rredlist (Gearty and Chamberlain, 2025) package to generate ranked lists automatically using the IUCN Red List API.

#### 2.3.1 Priority 1

Priority 1 species are evaluated against four conditions: they must be at risk of extinction, must be declining, must have a restricted population or range, and they must occur in the focal country, with those species with a higher degree of endemism scoring highest. Each of these conditions is quantified as described below.

***Risk*** weights the species according to their global IUCN Red List category on a scale of 0 to 4 (Table 1).

**Table 1.** Risk weights: Weights are selected to ensure that species are sorted by extinction risk category. See 2.3.3 for remark on EW.

| **Weight** | **Red List category** |
|---|---|
| 4 | CR |
| 2 | EN |
| 1 | VU |
| 0 | otherwise (LC, NT, DD, EX) |

***Decline*** weights the species according to urgency of conservation actions needed, approximated by the rate of population decline (Table 2). It is based mostly on IUCN Red List criteria A, C1 and E (IUCN, 2012). However, when available in the IUCN Red List assessment, data on the population trend and whether the species is coded as undergoing a ‘continuing decline’ are also used because a declining species may not meet IUCN Red List criteria A or C1 (e.g., a species listed as CR D alone may be declining at an inferred/suspected rate of 30-79% over 3 generations). To generate the weights, we first sorted the combinations of these metrics from less to more rapid declines, and then gave them values from 1 to 10 (see Table 2). In addition, we assigned a weight of 0.5 to unknown population trends to prevent the exclusion of species in decline although not known to be doing so, while accepting that many species not actually in decline will also be included. All other cases were assigned 0.

**Table 2.**
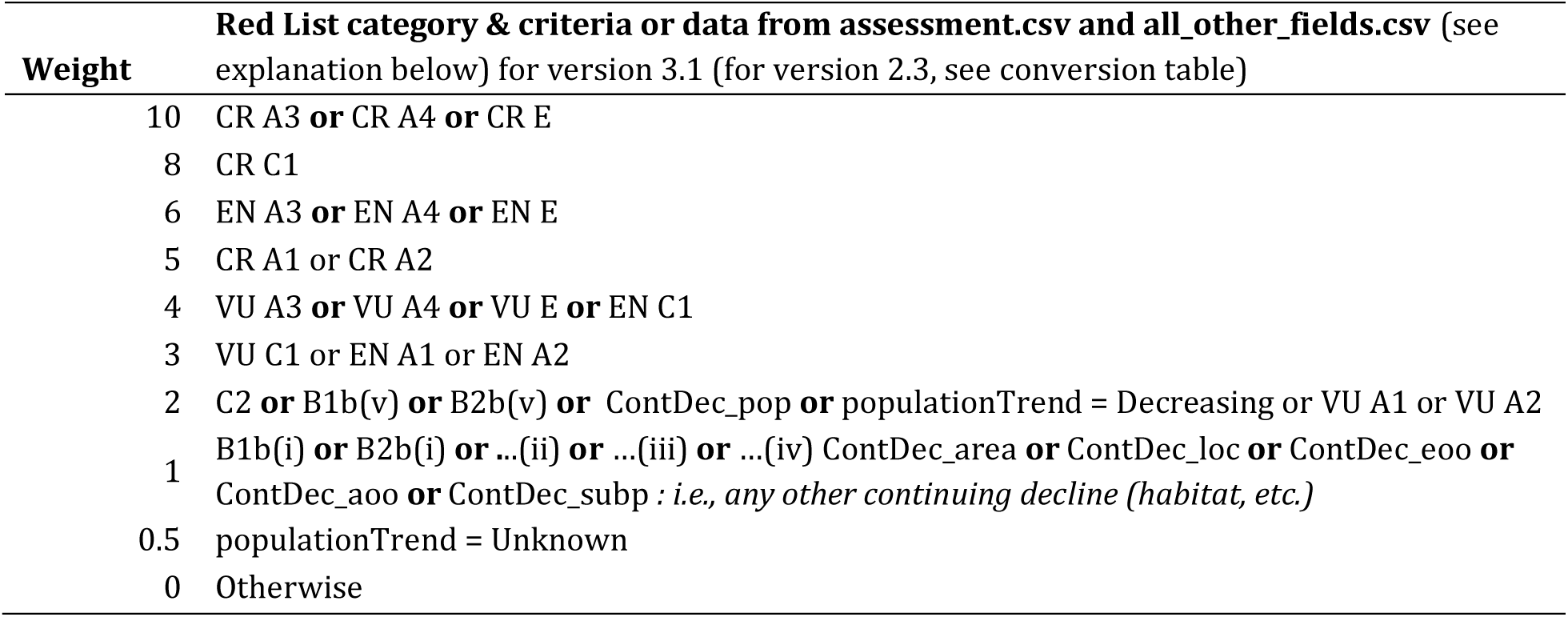
Decline: Reduction (ongoing or future) or continuing decline or population trend. Select the highest weight that the species qualifies for. The weights are based on the listing criteria and *populationTrend*, which is information not used in determining the Red List category, but is useful for detecting decline that the listing criteria do not include (e.g., a species that is listed as CR D but would have also met VU A2 or B2b(v)).

***Restriction*** weights the species according to the size of population or range (Table 3). It is based mostly on IUCN Red List criteria B, C, and D. However, as with Decline, other supporting data fields (e.g., population size, locations) from the IUCN Red List assessment are also used because a restricted species may not be listed with criteria B, C, or D (e.g., a species listed as CR A2 may have fewer than 250 mature individuals). Some metrics (such as population size) measure restriction directly while others (such as number of locations) imply restriction or measure it indirectly. To generate the weights, we first sorted the combinations of the metrics related to population size or range area from less to more restriction, and then gave them values from 1 to 10 (Table 3). All other cases were assigned 0.

**Table 3.** Restriction: Small population size or restricted distribution. Select the highest weight that the species qualifies for. The weights are based on the listing criteria as well as metrics indicating restriction that may not be reflected in the listing criteria (e.g., a species listed as CR A, but would have also met VU D or B1).

| **Weight** | **Red List category & criteria or data from assessment.csv and all_other_fields.csv**<br>(see explanation below) for version 3.1 (for version 2.3, see conversion table) |
|---|---|
| 10 | CR D **or** PopulationSize<50 |
| 9 | CR B **or** AOO<10 km<sup>2</sup> **or** EOO<100 km<sup>2</sup> **or** PopulationSize<100 |
| 8 | CR C **or** EN D **or** PopulationSize<250 |
| 7 | PopulationSize<500 |
| 6 | VU D1 **or** PopulationSize<1000 **or** LocationNumber = 1 |
| 5 | EN C **or** PopulationSize<**2500 or** LocationNumber = 2 |
| 4 | LocationNumber ≤ 5 **or** PopulationSize<5000 **or** VU D2 **or** AreaRestricted |
| 3 | EN B **or** AOO<500 km<sup>2</sup> **or** EOO<5000 km<sup>2</sup> |
| 2 | VU C **or** PopulationSize<10000 **or** LocationNumber ≤ 10 |
| 1 | VU B **or** AOO<2000 km<sup>2</sup> **or** EOO<20000 km<sup>2</sup> |
| 0 | Otherwise |

***Endemism*** weights species according to the number of countries the species occurs in, using this as a proxy for the proportion of the species’ distribution that the country comprises (Table 4). Species which are present (‘Extant’ or ‘Possibly Extinct’ if threatened), or have occurred in the focal country prior to extinction (‘Extinct Post-1500’ if EW) with native origin were selected. We assigned a value of 10 to extant species found only in one country (i.e. endemic), and assigned smaller weights to species found in more countries (Table 4). In all other cases Endemism is calculated to be ten divided by the number of countries in which the species occurs.

**Table 4.** Endemism: The importance of the country for the conservation of species, based on the number of countries that the species occurs in.

| **Weight** | **Condition** |
|---|---|
| 10/(number of countries in which the species is extant) | If the species is extant ( and known to be native, or believed plausibly to be so) in the focal country |
| 0 | Otherwise |

A priority score (PS1) based on these 4 weights is calculated using the following equation, and used to rank species:

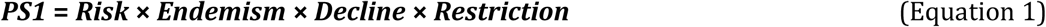

A species must thus score >0 for each of the four conditions to be included in the first part of the list (Priority 1). Species with *PS1*>0 are ranked from highest priority score to lowest. Species with *PS1*=0 are considered for the second part of the list (Priority 2, see 2.3.2). It is important to note that the priority score values are not necessarily linear functions of any particular metric of importance or urgency of action (e.g., *PS1*=2000 cannot be interpreted to be twice as important as *PS1*=1000). The priority score is primarily used for ranking only but given the standardised approach it could be used to compare between countries’ lists.

#### 2.3.2. Priority 2

For species with PS1=0, further consideration is necessary to ensure that some species are not excluded simply owing to data inadequacy or uncertainty. As Equation 1 does not consider restricted species to be a priority unless they are also declining, and does not consider declining species to be a priority unless they are also restricted, a second score (PS2) is calculated, to rank species that are only declining or only restricted. It also includes the species for which Decline or Restriction data may not be available. For example, a species may be listed as CR A2 on the IUCN Red List, and may also have a small population (i.e., would have met EN D or VU D), but the assessors may not know this or it may not be documented in the Red List assessment because the population size is not small enough to qualify the species as CR under criterion D. Or, a species may qualify as CR under criterion D, and be declining, but the assessors may not know the rate of decline, or may not record this in the assessment if it is not sufficient to meet criterion A at the CR level.

The above process is repeated using a modified equation to generate a second priority score (PS2):

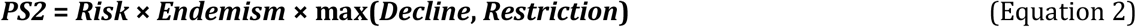

Equation 2 adds species that are not declining (but are restricted), and species that are not restricted but are declining, to the list. For example, a species listed as CR under criterion D with an increasing or stable (or unspecified) population trend would not be included on the list under Priority 1, but would appear in the Priority 2 part. Similarly, a species listed as CR under criterion A with a population size greater than 10,000 (or unspecified) would not be included under Priority 1 but would be included under Priority 2. PS2 is calculated only for species with PS1=0.

Species with PS2>0 are listed below species with PS1>0 to create a single ranked list.

#### 2.3.3. Extinct in the Wild species

Species which are listed as EW on the IUCN Red List were not evaluated against the four conditions and were instead placed unranked at the top of the combined list created from the two processes described above. This was done to highlight EW species and their complete dependence on ex-situ conservation (as called for in Target 4), while recognizing, too, their potential for recovery. However, we do not rank them, so that weavoid making a judgement on their priority relative to other globally threatened species.

### 2.4 Pilot testing

Participants from eight countries that are Parties to the CBD were approached to pilot test the above protocol, to assist them in addressing Target 4 of the KMGBF: Colombia, Fiji, Greece, Indonesia, Nigeria, Papua New Guinea, the Seychelles, and South Africa. These countries were selected to represent a range of contexts, varying in geographic region, latitude, species richness, and wealth, and included both continental and island states. Participants in each country came from a range of organisations including government departments, research institutes, non-governmental organisations, and IUCN SSC National Species Specialist Groups (Table 5), all of which are involved in national decisions concerning species and the KMGBF.

**Table 5.**
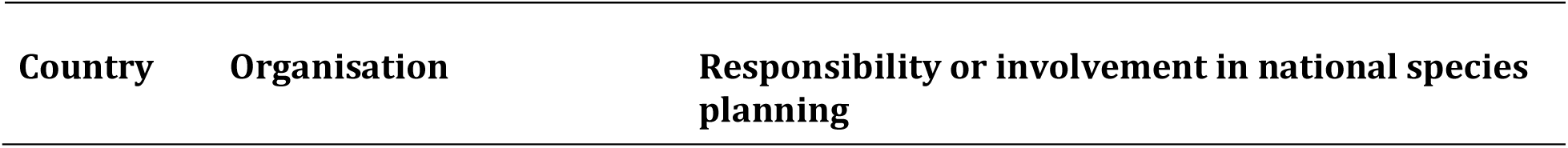
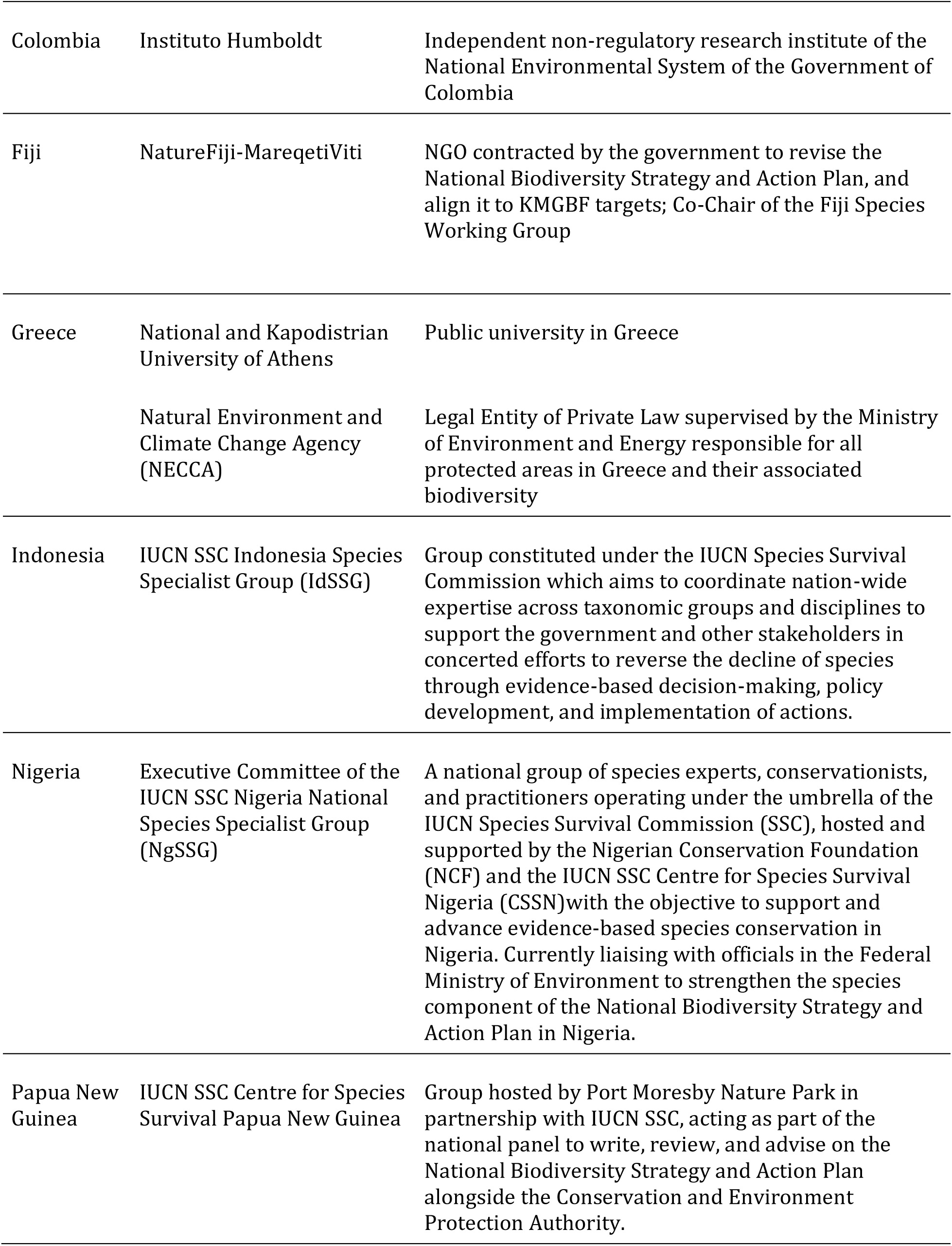
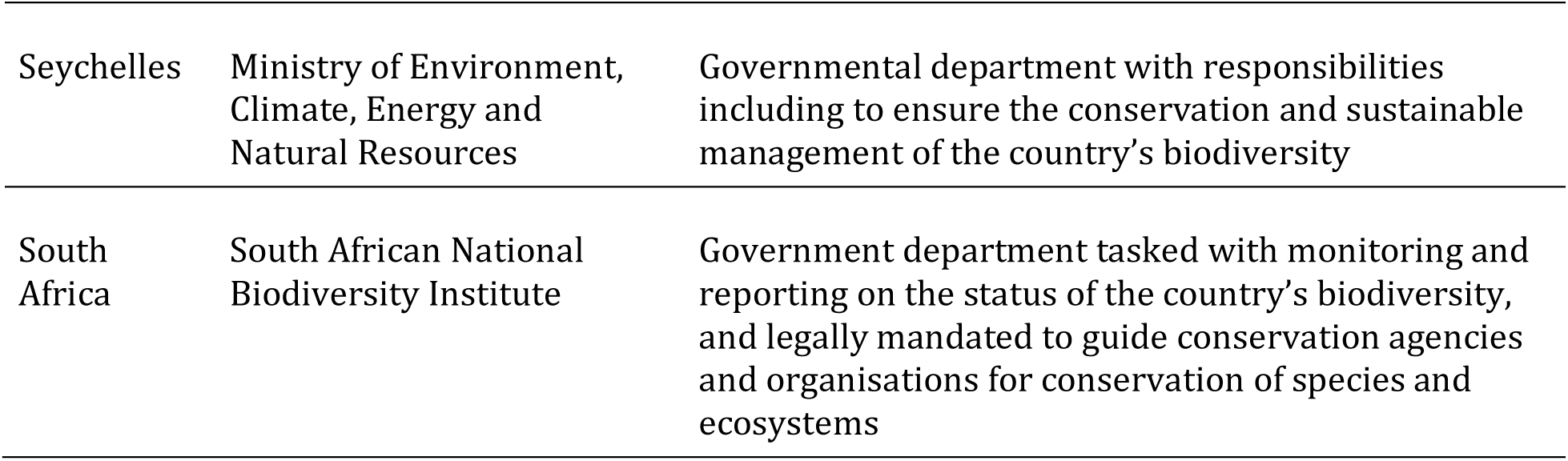
Summary of the organisations acting as country participants involved in the development and testing of this protocol.

A ranked list of species potentially requiring urgent management action for each country, identified using the protocol above, was shared with representatives of each organisation. We then asked each participant organisation whether the protocol is useful to them in achieving Target 4, and for comments on the value of the approach. We also inquired about potential next steps and support needed to finalise a list of species in need of urgent management action and to act on this list. The feedback was obtained using an online survey including a mix of open and multiple-choice questions (see Supplementary material 1), and from ongoing communication with representatives of the organisations listed in Table 5. We requested one representative response from each organisation, produced by internal consultation.

## 3. Results

### 3.1 Summary of ranked lists for all countries globally

Across all 250 countries and territories present in the IUCN Red List country list (IUCN, 2025, Brummitt, 2001), the number of species identified under P1 ranged from zero to 3,746 with a mean of 220 (SD 458; Fig. 1a). Five countries had over 2,000 species identified under P1, these were Madagascar (3746), Brazil (2403), Mexico (2296), Ecuador (2148) and Indonesia (2096).

**Figure 1.**
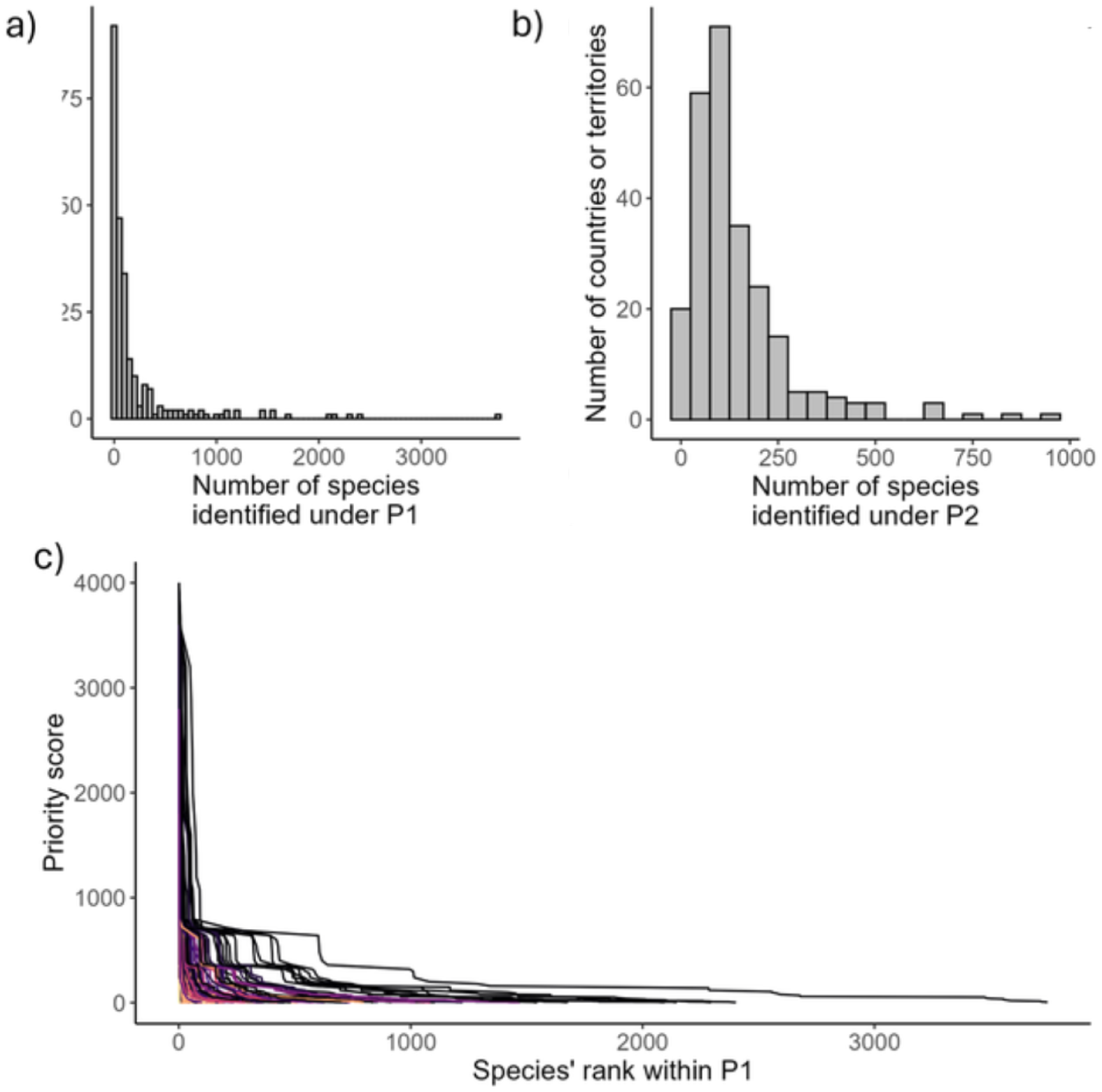
Summary of priority lists across all 250 countries & territories (defined using the [IUCN country codes list](https://www.iucnredlist.org/resources/country-codes)) a) the number of species identified under P1, b) the number of species identified under P2 and c) the relationship between national rank and priority score for species identified under P1 where low values for rank indicate higher priorities. Line colours indicate the priority score of the species ranked first (i.e. highest) for that country or territory with dark colours indicating high scores and light colours indicating low scores. This figure (a-c) excludes EW species which are unranked.

The lowest priority score of any species identified under P1 (i.e. the score of the species ranked last in P1) ranged from 0.38 - 80 among countries, with a mean of 2.4 (SD 6.50). The priority score of the highest ranked species ranged from 0.87 - 4000 among countries and territories, with a mean of 1375 (SD 1321.75). Twenty countries had at least one species with the highest possible score for a species extant in the wild of 4000 (45 species in total; Fig. 1c).

Four countries or territories had no species identified under P1 (Guernsey, Holy See, Maldives and San Marino) and a further seven had one species identified (Åland Islands, Bouvet Island, an un-named Disputed Territory, Nauru, Saint Pierre and Miquelon, Svalbard and Jan Mayen and Tokelau). Among all 85 countries or territories that had 20 or fewer species identified under P1, the number of species identified under P2 ranged from 3 to 283 with a mean of 82 (SD 62). Across all 250 countries or territories, the number of species identified under P2 ranged from 3 to 944 with a mean of 143 (SD 139; Fig. 1b).

Among the 40 countries associated with at least one EW species, the number of EW species listed ranged from one to twelve. These species were listed as a priority but not ranked.

### 3.2 Ranked lists for the eight pilot countries

The proportion of Priority 1 species included in national ranked lists produced by our protocol ranged between 52.7% and 86.1% of globally threatened species for our eight pilot countries (Table 6). For all eight countries, excluding Nigeria, the majority of Priority 1 species were endemic (Fig. 2A). The proportion of species within each of the IUCN Red List categories varied between countries (Fig. 2B). Taxonomic representation differed between countries’ ranked lists, but Tracheophyta, Arthropoda, Chordata, and Mollusca were present in all (Fig. 3). Figures showing the distribution of species across endemism and extinction risk categories are shown in Supplementary material 2.

**Figure 2.**
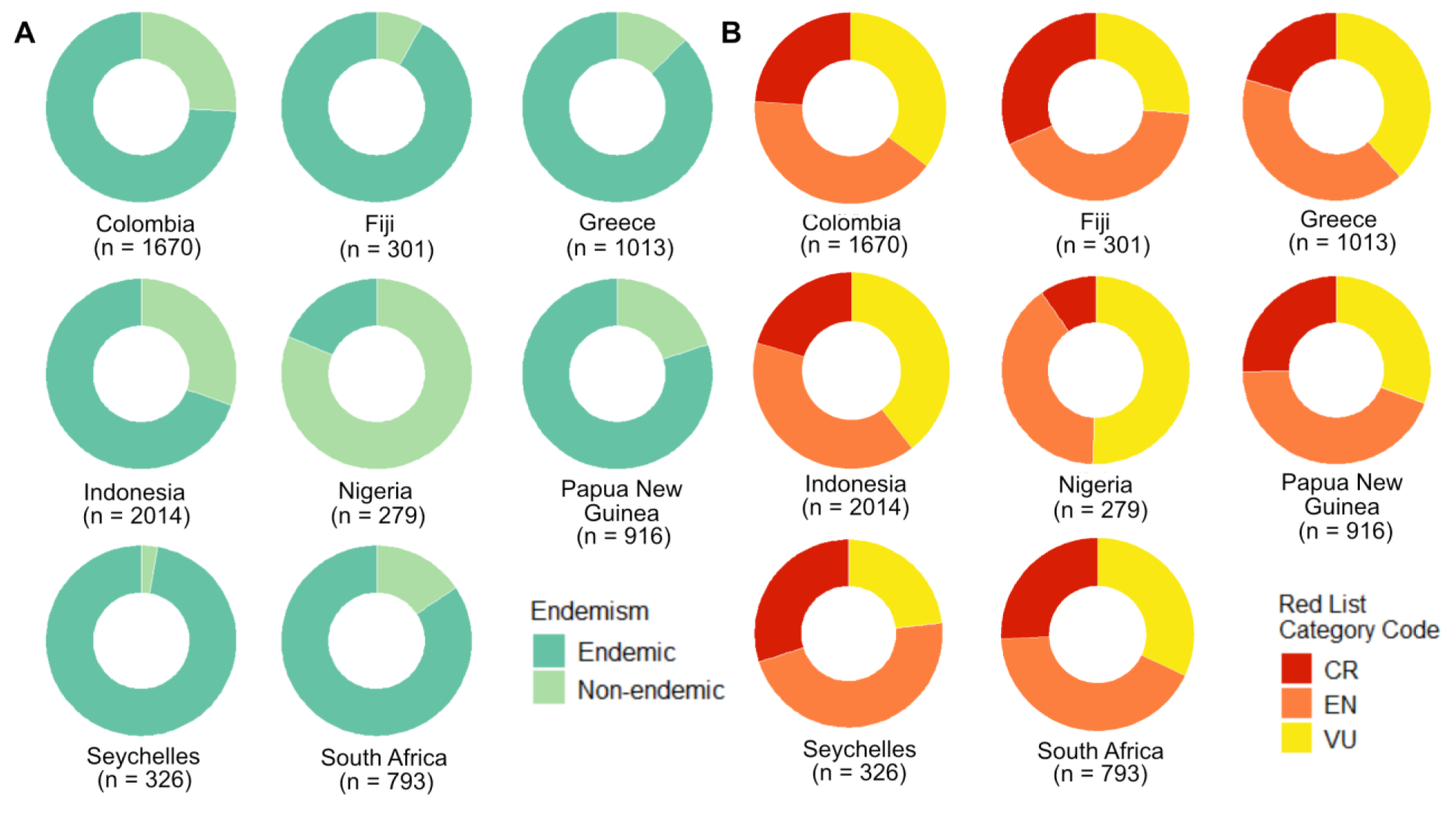
The proportion of species included on the Priority 1 part of the ranked lists produced by the protocol A) that are endemic to each of the eight pilot countries, and B) that fall within each of the IUCN Red List categories. The number of Priority 1 species for each country is indicated in parentheses after the country name. EW species are excluded from this figure.

**Figure 3.**
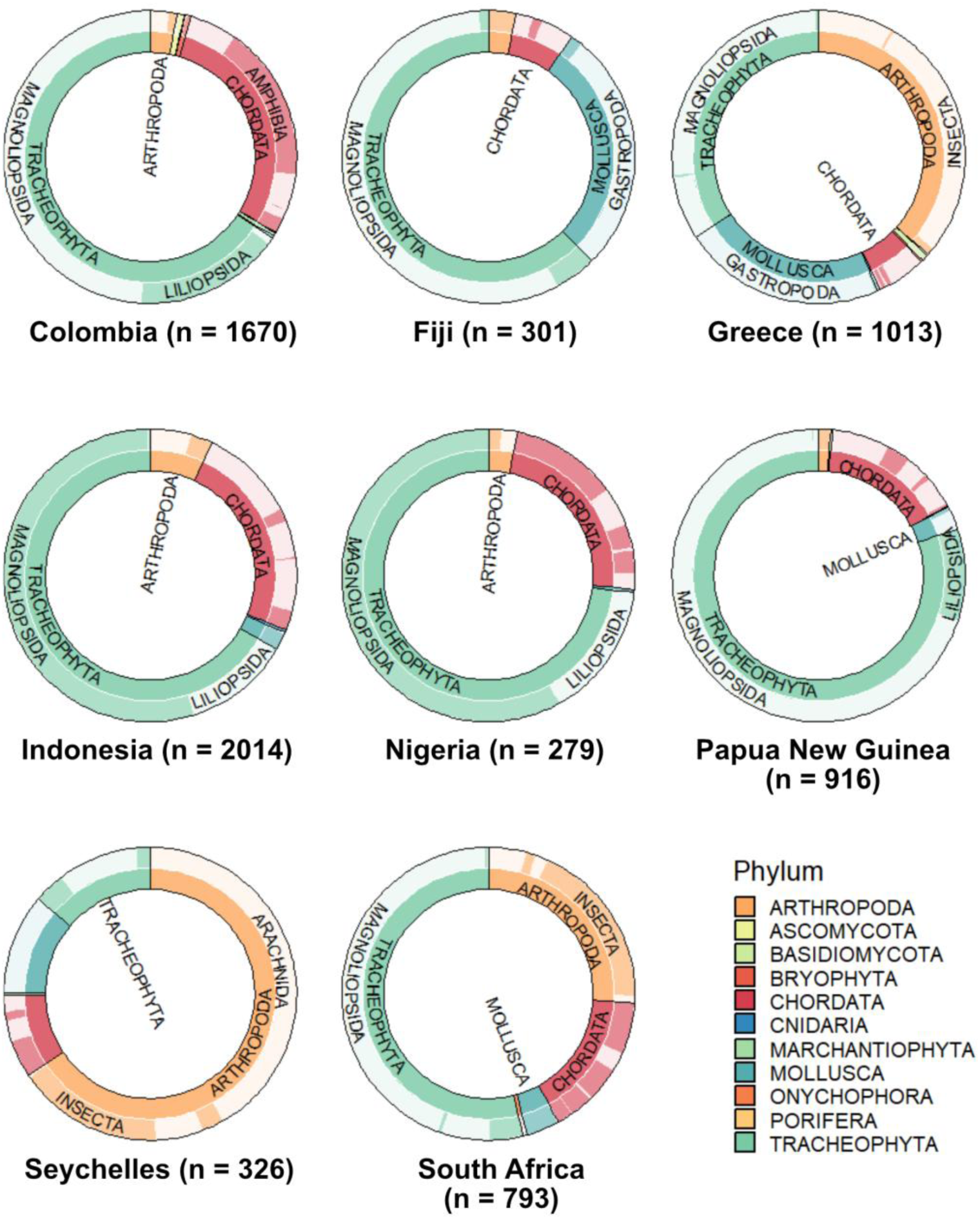
The proportion of species in each phylum (inner circle) and class (outer circle) included on the ranked list of Priority 1 species for each of the eight pilot countries. The total number of Priority 1 for each country is indicated in parentheses after the country name. This figure excludes EW species.

**Table 6.** Summary of the number of threatened species listed on the IUCN Red List occurring within each of the eight pilot countries, the number of species in the Priority 1 section of the list produced by this protocol, and the proportion of threatened species represented by Priority 1 species for each pilot country.

| **Country** | **Number of threatened and EW species on IUCN Red List** | **Number of species identified under Priority 1 on the ranked list produced by this protocol** | **Proportion of threatened species represented by Priority 1 species** |
|---|---|---|---|
| Colombia | 2,002 | 1,670 | 83.4% |
| Fiji | 571 | 301 | 52.7% |
| Greece | 1,176 | 1,013 | 86.1% |
| Indonesia | 2,919 | 2,014 | 68.9% |
| Nigeria | 510 | 279 | 54.7% |
| Papua New Guinea | 1,322 | 916 | 69.3% |
| Seychelles | 563 | 326 | 57.9% |
| South Africa | 1,086 | 793 | 73.0% |

### 3.3 Pilot testing: Participants’ perceptions of the ranked lists for urgent management action

Participants were presented with a national ranked list for their country, produced using the protocol described above, and given the opportunity to review them prior to answering these questions.

#### 3.3.1 Objective 1: Is this protocol useful?

All the participating groups agreed that a ranked list produced by a standardised protocol such as this one would be helpful in addressing Target 4 (Fig. 4). This was considered to be particularly helpful in directing action to the species most in need, saving time in identifying species, providing a starting point from which a national priority list of species can be identified, and creating confidence that the process of selecting species was scientifically robust. Additional reasons the list was considered helpful was for reducing bias, and highlighting species in need of focused in-country research.

**Figure 4.**
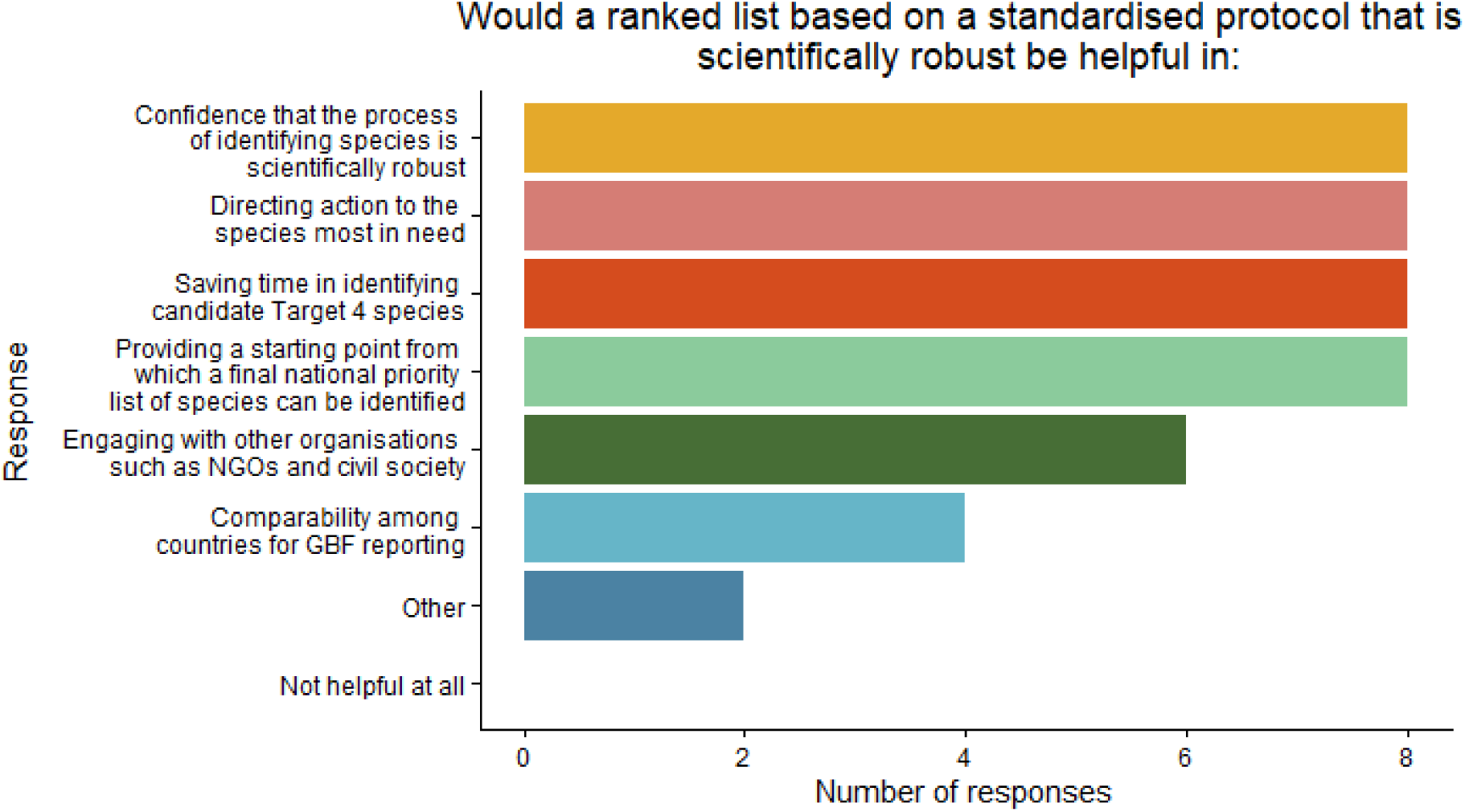
Multiple choice responses from eight countries to the question “Would a ranked list based on a standardised protocol that is scientifically robust be helpful in…”.

#### 3.3.2 Objective 2: What are the next steps needed to finalise a national list of species in need of urgent management action?

Participants were asked about potential next steps they would undertake to create a national list of species in need of urgent management actions, using our ranked list as a starting point. This was to better understand where our protocol sits within the political decision-making process of countries (e.g., influencing national biodiversity strategy and action plans).

Participants were asked whether having a shorter list of species that are absolute priorities to be addressed under Target 4 would be helpful, reflecting prior conversations with in-country partners indicating this would be more practical for selecting species in need of urgent management actions. Participants from Colombia and South Africa indicated that they would not find this helpful, as they have mechanisms in place that could be used to shorten the full ranked list and prioritise species, e.g. by consulting with stakeholders and species experts. Participants from the remaining pilot countries indicated that they would find such a shorter list of species helpful, but do not yet have a process in place to implement this. These six countries varied in considering which criteria should be used to create such a shorter list (Fig. 5) but unanimously considered specific biological criteria (e.g. endemism, population trend, overall population size or biological criteria not already included in the priority score) to be important in refining ranked species lists.

**Figure 5.**
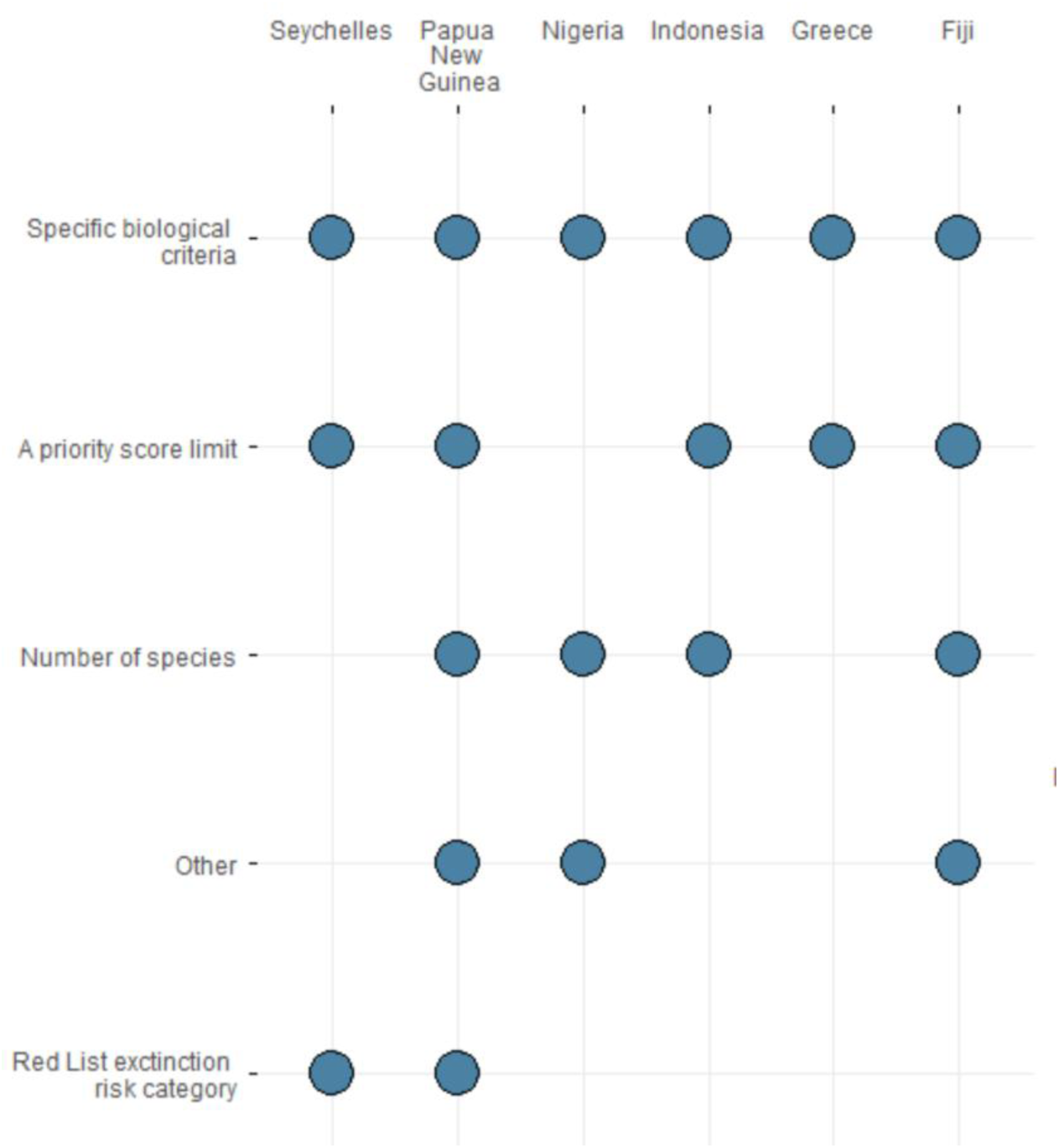
Balloon plot summarising the multiple-choice answer to “If it would be helpful to have a shorter list of species that are absolute priorities to be addressed under Target 4, how should this shorter list be defined?” (n = 6).

Participants from all eight countries welcomed the list, but felt some modifications should be made to better suit national contexts. The protocol’s selection criteria were chosen to reflect the wording of Target 4, but Parties may wish to apply additional criteria for sub-prioritisation of species for targeted urgent management actions. The essential criteria that participants indicated were important to consider, in addition to risk of extinction used to produce the ranked list, differed substantially between countries, with each considering a different combination of factors a priority (Fig. 6). Suggested next steps (given as open responses) included approaching species experts for advice and feedback on the ranked lists e.g. ensuring the national list was representative across taxa and habitats, assessing current species-level conservation efforts, determining an appropriate cut-off within the ranked list or otherwise producing a shorter list of species, and assessing the feasibility and costs of recovery actions.

**Figure 6.**
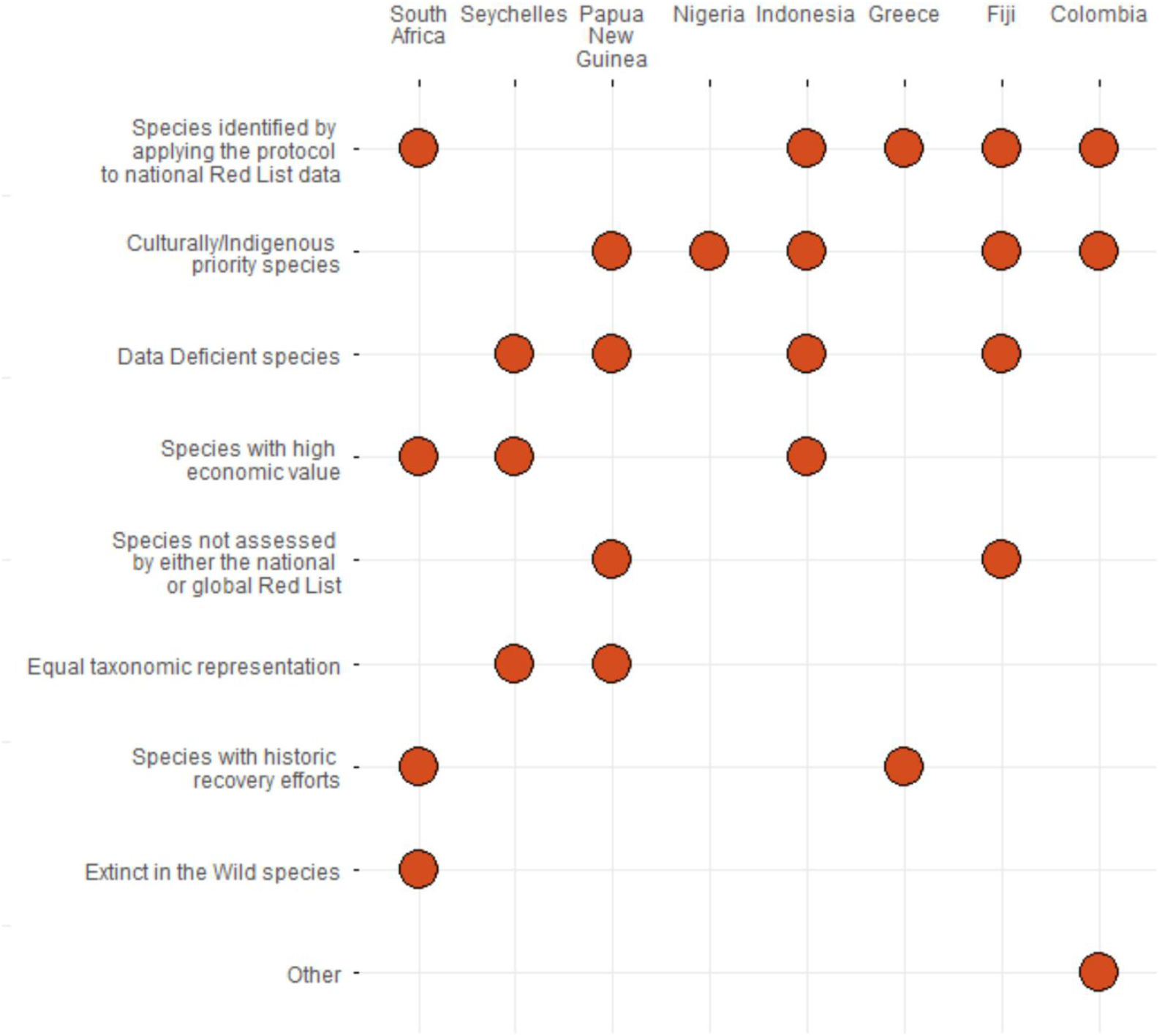
Balloon plot summarising the multiple-choice answer to “The protocol uses four criteria to prioritise species. Which (if any) of the following additional criteria are essential to consider when prioritising species within your national context?” (n = 8).

Participants anticipated hurdles to the finalisation and adoption of a national list of species in need of urgent management action under Target 4. Open responses included time (e.g. to obtain expert guidance, to update information, to develop and adopt the list), lack of funding, resources, or expertise, and engaging collaborators.

#### 3.3.3 Objective 3: Is further support, in addition to the ranked list, needed to act on this list?

Parties described a wide variety of additional support they would require in order to act on the list produced by our protocol, both to adopt a national Target 4 species list and to implement recovery-focused actions for the priority species. The required resources and guidance varied across the diverse national contexts considered here (see Table 7), but included guidance on developing species recovery plans, training and guidance on using the protocol, technical expertise on undertaking national Red List assessments, and production of a website with information on the priority species.

**Table 7.** Summary of country participants’ responses detailing what additional guidance and tools they would need to adopt a national Target 4 species list, and undertake recovery actions for these species.

| Country | What additional guidance or tools would help support you to adopt a national Target 4 species list? | What additional guidance or tools would help support you to mobilise recovery-focused actions for the identified species? |
|---|---|---|
| Colombia | Access to additional species information (e.g. specific threats, geographical data) to refine the prioritisation of the list at the national level | Background on recovery actions that could be implemented and case studies with lessons learned, particularly for groups that have received less attention e.g., insects, bryophytes, and fungi |
| Fiji | Training on using the protocol, Target 4 process, and IUCN Red List assessments for national species specialists so that the knowledge holders (both western and traditional) can agree to a national Target 4 species list | Training for conservation action planning and fundraising for species conservation action<br>Funds to implement agreed actions<br>An accessible online toolkit with self-help material related to Target 4, a communications strategy to create awareness around Target 4 species, a dedicated webpage with information on Fiji’s Target 4 species |
| Greece | Adoption of a national Target 4 list requires political support for a science-based list | Identifying species experts, recommended conservation measures for the same or similar species |
| Indonesia | Clear guidance on how the ranked list should be interpreted and adapted nationally, including minimum standards for validation and refinement<br>Technical tools or dashboards, enabling easier filtering, updating, and visualisation of species by threat level, geography, and management need<br>Guidance on governance and ownership, clarifying how national institutions can formally adopt, maintain, and report on the list over time | Action-oriented guidance, linking priority species categories to recommended types of recovery interventions (e.g. habitat protection, threat mitigation, population management) |
| Nigeria | Technical and policy advocacy support for future engagement | Support and access to training on species management and recovery plan development, and capacity development for implementation |
| Papua New Guinea | Analysing the species list against Key Biodiversity Areas and protected areas | Linking resources with in-country institutions and NGOs, training for stakeholder mapping, analysis, and species assessments, funding |
| Seychelles | Technical expertise, capacity building and training on developing a national Red List | Data collection guidance, monitoring protocols, training materials on undertaking biodiversity assessments |
| South Africa | A guidance document to help countries select Target 4 species | A website advertising the species in need of recovery, including clearly identified actions and costs of recovery |

## 4. Discussion

Our novel protocol provides a standardised method of identifying species potentially in need of urgent management actions to prevent their extinction and enable their recovery. While there are limitations to the protocol, which we consider below, by producing ranked species lists we provide a critical starting point to support Parties in identifying national priority species, and addressing Target 4 of the KMGBF. The output was found helpful by participants from relevant institutions or groups in the eight countries in which it was piloted. Our results also highlight the need for context-specific next steps and additional support to develop and act on national priority lists.

Participants who piloted this protocol agreed unanimously that it was helpful. The ranked lists save time in identifying species, which is important given the limited time available to make progress on achieving Target 4 before 2030. Participants also supported the protocol due to it providing a starting point from which to identify a national priority list of species, and directing action to the species most in need, substantial challenges which contributed to Parties not achieving Aichi Biodiversity Target 12 (CBD, 2016b). Application of this protocol can assist in addressing a substantial challenge through a scientifically robust process that can be applied for all countries, and in which Parties may have confidence.

### 4.1 Protocol evaluation

While the protocol provides a useful foundation for identifying species needing urgent management action, there are additional considerations not captured by our approach. The IUCN Red List also includes information on threats, stresses, and conservation actions, which can be appropriate for broad-scale assessments (e.g., Bolam et al. 2023). We chose not to use these data here due to some documentation being incomplete or not required, which could result in inaccuracies when ranking species. Because threats and required actions vary by context and location, expert evaluation is likely required to determine whether species-targeted measures are appropriate or feasible within a given country. Thus, countries should adjust the priority list by removing species already well addressed by actions under other CBD targets, adding species that require targeted national action, or downgrading species unlikely to benefit from urgent management within their jurisdiction. The Species Threat Abatement and Restoration metric (Mair et al. 2021) can be used to calculate the relative contribution of threats to overall extinction risk in a country, and indeed at any spatial scale, and this will inform which GBF targets would contribute the most to the reduction of species extinction risk, and which species those targets cover.

To apply the Endemism criteria we used a proxy measure of the proportion of countries/territories (as defined by IUCN) occupied by a species that is represented by the focal country (Table 2). The proportion of a species’ range or Area of Habitat (AOH; Brooks et al 2019) in the focal country would provide a more accurate proxy of each country’s importance for the conservation of a species (i.e. the proportion of the global population it supports) where these data are available. For example, using the current protocol one species with 50% of its range in Country A and 50% in Country B, and another species with 90% of its range in Country A and 10% in Country B, would both receive a score of 5, which may underestimate country A’s importance for the second species. However, as range maps and AOH maps are not yet available for all species assessed on the Red List, we chose to use the described method to reduce bias in species rankings. Maps for all species assessed on the Red List are expected in the near future. Spatial data on species’ distributions are now required for all updated Red List assessments, so as older assessments lacking such documentation are updated, it will be possible to calculate endemism scores more accurately.

The protocol uses data from all species listed on the IUCN Red List, and is not limited to comprehensively assessed groups (i.e. those in which >80% of species in the group have been assessed). This was done to reduce taxonomic bias and address the challenge of charismatic species disproportionately receiving conservation attention (Guénard *et al*., 2025). Some country participants, including from Greece and Seychelles, highlighted that this impartiality allowed some non-charismatic species to rank highly, which they felt was a benefit of the protocol, as the species may otherwise be overlooked. However, there is uneven assessment coverage across different taxonomic groups within the IUCN Red List. For example, all tetrapods have been comprehensively assessed, versus 18% of described plants and <5% of fungi (IUCN, 2025). Certain countries have better representation across taxonomic groups included in national red list assessments (Raimondo et al. 2022). For countries utilising the IUCN 3.1. Red List criteria for national assessments, such as Colombia (Rodríguez et al. 2006), Greece (NECCA, 2024), and South Africa (https://speciesstatus.sanbi.org/), we encourage that this same prioritisation script be applied to species present on national red lists but absent from the global red list, in particular national endemics, to augment the Target 4 listings.

Furthermore, the proportion of fishes, invertebrates, plants, and fungi with Red List assessments differs between countries, which influences how highly particular assessed species in these groups and among tetrapods (which are comprehensively assessed) appear in the ranked lists, and hence potentially which are prioritised for urgent management action. We included all assessed species in each country because Target 4 refers to “species” and “known threatened species” in general. Nevertheless, it is important to be aware that the priority lists produced using this protocol reflect the taxonomic and geographic composition of the underlying data. Ranked lists will also require updating as species are reassessed and added to the Red List.

There are also rare cases where some species may not be ranked highly by our protocol, but should be prioritised for urgent management action in a particular country. For example, there may be cases where Endemism as applied here is a poor proxy for the proportion of the global population occurring in a country and hence the nation’s importance for a species’ conservation, such as migratory species dependent on ‘bottleneck’ sites within the country. In addition, our protocol excludes Data Deficient species, which may require targeted conservation. Our pilot study also identified a small number of endemic species with very restricted ranges within a country, which had equal ranking with other endemic species that occur throughout the country, such as *Hypogeophis montanus*, a caecilian which occurs only on the top of two mountains on Mahé in Seychelles (Maddock, Wilkinson and Gower, 2018), compared with e.g., *Nesiergus gardineri*, which is found on multiple islands in Seychelles. The ranking of such species may be refined with specialist technical input (e.g., by breaking tied ranks using additional criteria such as the number of occupied islands).

Finally, it is worth highlighting that incomplete Red List supporting documentation (e.g. criteria incorrectly coded up), or errors, may skew or influence results. Age of assessment should also be considered (more than one-fifth of Red List assessments are currently marked as needs updating, i.e. over 10 years old); while the data may still be relevant, it is also possible that newer data may be available to inform the ranking process. These situations illustrate that, while the protocol is useful for identifying a list of species that would benefit most from urgent management actions, the ranked lists should be manually reviewed and revised accordingly.

### 4.2 Next steps

Ideally, countries would select and take action to protect all threatened and EW species within their borders. Realistically, however, many countries need to prioritise those species that require the most urgent management actions (highly ranked species), with other species being added as resources and capacity become available. Attempting to take action for all threatened, or all Priority 1 species within a country may spread resources too thin and result in insufficient action for the species closest to extinction. National refinement of the ranked lists will be necessary, as there is no one-size-fits-all method for developing priority species lists; countries differ in context and approaches, as shown in our pilot testing. For example, Nigeria refined its list by selecting a target number of endemic species and adding nationally important species from its NBSAP. Each pilot country identified its own criteria for narrowing priorities, including national Red List data, culturally significant or Indigenous priority species, and Data Deficient species, indicating that the most appropriate method will be country-specific. This aligns with Consideration D of the KMGBF (CBD, 2022a), which states: “Each Party would contribute to attaining the goals and targets of the Framework in accordance with national circumstances, priorities and capabilities”. As many species requiring urgent management action under Target 4 occur in more than one country, agreement on shared prioritisation among neighbouring countries may lead to increased conservation impact without increased effort. The CBD’s Technical and Scientific Cooperation Centres (CBD 2022b; https://www.cbd.int/tsc/tscm/subregionalcentres) could play a key role in facilitating such discussions and applying the protocol across borders to maximise the impact of urgent management actions taken nationally.

On capabilities, Parties face numerous other barriers to achieving Target 4, which this protocol does not address. In some pilot countries, the number of species identified as needing urgent management exceeded their economic capacity. This is a similar challenge to area-based conservation targets (Shen *et al*., 2023) and may particularly impact small island developing states with relatively high levels of endemism like Fiji and Seychelles. A priority list may serve to catalyse efforts to increase funding for species conservation in such cases. The implementation of conservation policy targets could be more efficient through effective prioritisation, for which we provide a solution here, but also if they take greater account of socioeconomic and cultural contexts, as human, institutional and financial capacities are critical to the overall ability of nations to address these targets (IPBES, 2019). There was substantial variation in responses from participant countries about the next steps, hurdles, and support required to implement and act on a national priority list of species for urgent management action based on our protocol. However, from these responses it is clear that additional guidance, training, and resources are needed to achieve meaningful progress on Target 4.

### 4.3 Scaling up

This protocol and resulting lists could inform all Parties in selecting species in need of urgent management action under Target 4. The benefits of this would be a standardised, scientifically robust, global starting point to approach selecting species, with greater comparability among KMGBF Parties. This approach is equally useful for the Convention on Migratory Species, the Ramsar Convention and other agreements aiming to address conservation of species. To this end, we provide the protocol code in an open-access repository (github.com/JonahMorreale/RedList_Target4Species). Reverse the Red is also building a set of support tools and frameworks to assist countries identifying and prioritising their Target 4 species, including requesting their national ranked list, which can be found on their [website](https://www.reversethered.org/). To support Parties, socialisation through strategic partnerships and international conservation networks, such as the CBD’s Technical and Scientific Cooperation Centres (TSCCs), IUCN Species Survival Commission, and the [Reverse the Red](http://www.reversethered.org/) coalition will be undertaken.

Goal A of the KMGBF sets out the outcomes that need to be achieved by 2050; in order to achieve these, we need to implement the actions outlined in Targets 1 - 8. Because political and ecological timescales vary (Watts et al., 2020) and it may take time to see these outcomes (Piipponen-Doyle, Bolam and Mair, 2021), it is important to have appropriate, action-based indicators to show progress is being made. An indicator based on “the proportion of species in urgent need of recovery that have recovery plans in place and implementation actions underway for Target 4”, was proposed by African Parties at the 26th meeting of the CBD’s Subsidiary Body on Scientific, Technical and Technological Advice in May 2024 (CBD, 2024). An indicator that tracks urgent management actions taken for priority species is clearly needed and reporting against such an indicator would quantify progress has been made up to 2030. Once Parties have selected appropriate priority species, we recommend they establish clear baselines from which to assess progress. Although global Red List assessments provide an initial extinction-risk baseline, some countries may require updated national assessments. Conducting Green Status of Species assessments can offer a robust recovery baseline and targets, including forecasts of Conservation Gain and Conservation Dependence over the next decade. As Parties implement and record conservation actions, progress towards species recovery can then be monitored using indicators such as the Green Status Index of Species Recovery (Akçakaya et al., 2025), already being piloted by South Africa and Indonesia, and the Red List Index (Butchart et al., 2025).

### 4.4 Conclusions

We have developed a transparent, standardised, and scientifically robust protocol to support Parties in identifying species potentially in need of urgent management action under Target 4 of the Kunming–Montreal Global Biodiversity Framework. Feedback from eight pilot countries indicates that the protocol is practical, efficient, and useful in generating starting points for national prioritisation. Refinement at the national level is important, reflecting country-specific ecological, socioeconomic, and institutional contexts, and further steps such as capacity building and technical support will strengthen consistency and uptake. Despite acknowledged data gaps and biases within the IUCN Red List, it remains by far the most comprehensive relevant global dataset currently available. With 2030 rapidly approaching, time is of the essence to achieve all KMGBF Targets. Using the best available data through a clear, justifiable, and globally applicable method offers an immediate and practical pathway for accelerating meaningful progress towards Target 4.

## Data statement

The reproducible pipeline including data retrieval via the Red List API, all data processing and protocol code, is available in an open-access repository: github.com/JonahMorreale/RedList_Target4Species.

## Acknowledgements

NM and BL would like to acknowledge Re:wild and the Lyda Hill Foundation for their support.

## Author contributions: CRediT

**H. Resit Akçakaya:** Conceptualization; Methodology; Writing – Original Draft; Writing – Review & Editing. **Natasha Mannion:** Writing – Original Draft; Project Administration; Investigation; Writing – Review & Editing; Visualisation. **Jonah Morreale:** Software; Methodology; Writing – Review & Editing. **Domitilla Raimondo:** Conceptualization; Methodology; Investigation; Writing – Review & Editing. **Michael Hoffmann:** Conceptualization; Methodology; Writing – Original Draft; Writing – Review & Editing. **Stuart H. M. Butchart:** Conceptualization; Writing – Review & Editing. **Louise Mair:** Conceptualization; Writing - Original Draft; Writing – Review & Editing. **Francesca A. Ridley:** Conceptualization; Writing - Original Draft; Writing – Review & Editing; Visualisation. **Malin Rivers:** Methodology; Writing – Review & Editing. **Caitlin Brant:** Conceptualization; Writing – Review & Editing. **Michael Clifford:** Conceptualization; Writing – Review & Editing. **Megan Joyce:** Conceptualization; Writing – Review & Editing. **Kira Mileham:** Conceptualization; Project Administration. **Celia Nova Felicity:** Investigation; Writing – Review & Editing. **Mirza Kusrini:** Investigation. **Sunarto:** Investigation; Writing – Review & Editing. **Johnpaul Houston:** Investigation; Writing – Review & Editing. **Nunia Thomas:** Investigation; Writing – Review & Editing. **Simon T. Maddock:** Investigation; Writing – Review & Editing. **José F. González-Maya:** Investigation; Writing – Review & Editing. **Kostas Triantis:** Investigation; Writing – Review & Editing. **Dimitrios Vavylis:** Investigation; Writing – Review & Editing. **Konstantina Spiliopoulou:** Investigation. **Indira A. Gamatis:** Investigation; Writing – Review & Editing. **Bello A. Danmallam:** Investigation. **Samuel T. Ivande:** Investigation; Writing – Review & Editing. **Shiiwua A. Manu:** Investigation. **Stella Egbe:** Investigation. **Joseph D. Onoja:** Investigation. **Carolina Castellanos-Castro:** Investigation; Writing – Review & Editing. **Cristina Lopez-Gallego:** Investigation; Writing – Review & Editing. **Barney Long:** Conceptualization; Writing – Review & Editing. **Philip J. K. McGowan:** Conceptualization; Investigation; Writing – Original Draft; Writing – Review & Editing.

