## Supplements for "Identifying and ranking species that need urgent management action to achieve Target 4 of the Global Biodiversity Framework"

**Supplement 1:** Survey questions posed to participants from eight pilot countries.

| Question | Question format | Multiple choice options |
| --- | --- | --- |
| Which Party do you represent? | Open |  |
| Please provide a contact name | Open |  |
| Please provide an email address | Open |  |
| Would a ranked list based on a standardised protocol that is scientifically robust be helpful in: | Multiple choice | Providing a starting point from which a final national priority list of species can be identified; Saving time in identifying candidate Target 4 species; Confidence that the process of identifying species is scientifically robust; Directing action to the species most in need; Comparability among countries for GBF reporting; Engaging with other organisations such as NGOs and civil society; Not helpful at all; Other |
| From the perspective of your agency and in your national context, to what extent does this list represent species in need of urgent management action under Target 4? | Multiple choice | Strongly represents; Somewhat represents; Poorly represents; Doesn't represent at all; Unsure |
| The protocol ranks all species with available data. Would it be helpful to have a shorter list of species that are absolute priorities to be addressed under Target 4? | Multiple choice | Yes; No |
| If your answer to Q6 was 'No', why not? | Open |  |
| If your answer to Q6 was 'Yes', how should this shorter list be defined? | Multiple choice | A priority score limit: (e.g. all species achieving a priority score above a certain value). The priority score is a combination of risk, endemism, decline, and population or range restriction.; Red List extinction risk category (e.g. Endangered or above). If yes, above which category?; Specific biological criteria (e.g. endemism, population trend, overall population size or biological criteria not already included in the priority score). If yes, which criteria?; Number of species (e.g. top 50 – 300 species). If yes, what number of species is appropriate?; Other |
| The protocol uses four criteria to prioritise species. Which (if any) of the following additional criteria are essential to consider when prioritising species within your national context? | Multiple choice | Species identified by applying the protocol to national Red List data; Extinct in the Wild species; Data Deficient species; Species not assessed by either the national or global Red List; Culturally/Indigenous priority species; Species with historic recovery efforts; Species with high economic value; Equal taxonomic representation; None of the above; Other |
| If you were to use the initial ranked list, what are the steps you would need to take next? | Open |  |
| What hurdles do you anticipate in the development and adoption of a national list of species in need of urgent management action under Target 4? | Open |  |
| What additional guidance or tools would help support you to adopt a national Target 4 species list? | Open |  |
| What additional guidance or tools would help support you to mobilize recovery-focused actions for the identified species? | Open |  |

**Supplement 2**: Figure showing the distribution of species across endemism and extinction risk categories.

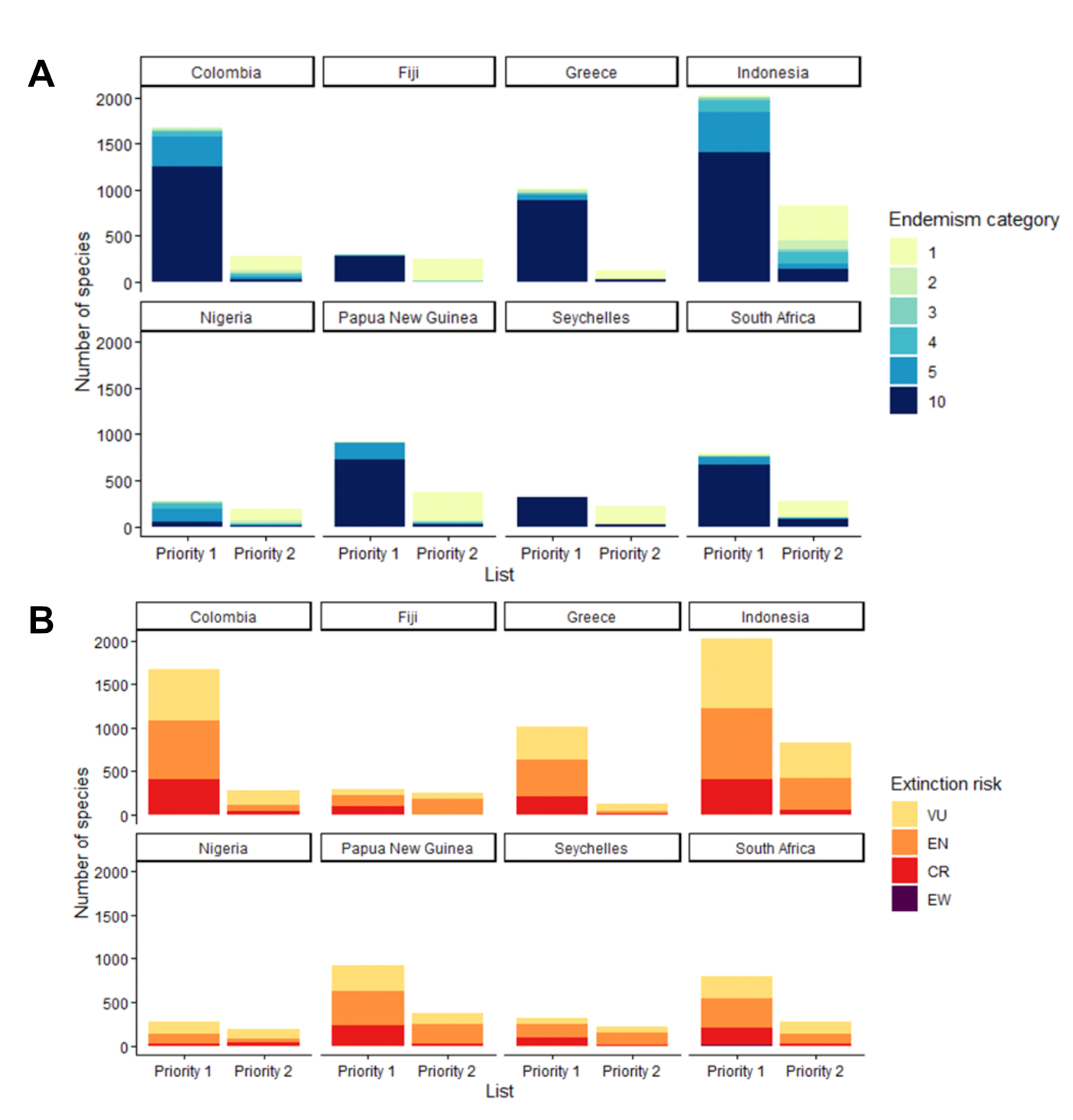

Figure S1. A) The number of species in each endemism category included on the ranked list of priority species for each of the eight pilot countries. B) The number of species in each IUCN Red List category included on the ranked list of priority species for each of the eight pilot countries.

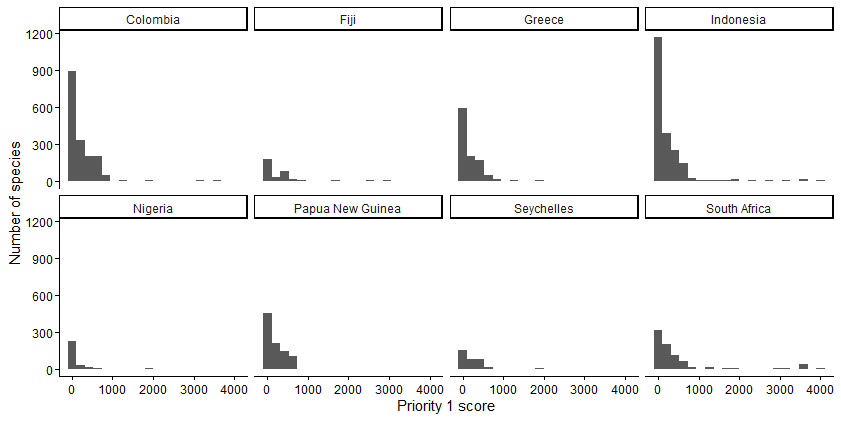

Figure S2. Histograms illustrating the frequency distribution of Priority 1 scores derived from the methodology proposed in this manuscript for the eight pilot countries considered.
